# Exploring the relationship between polymorphisms of leptin and IGF-1 genes with milk yield in indicine and taurine crossbred cows

**DOI:** 10.1101/814004

**Authors:** Mohammad Rayees Dar, Mahendra Singh, Sunita Thakur, Archana Verma

## Abstract

Leptin and IGF-1 plays a significant role in milk production and lactation in bovines. The present investigation was carried out to identify the novel polymorphisms in exon 3 region of leptin gene and exon 3 + partial intron 3 of IGF-1 gene and to analyze their association with the milk production performance in indicine and taurine crossbred (Karan Fries) cows. Blood samples were collected from 160 apparently healthy Karan Fries cows. Four SNPs at positions rs29004508 (C>T), rs29004509 (C>T), rs29004510 (T>C), rs29004511 (T>C) in Leptin gene and two SNPs at positions rs133251968 (C>A), rs137289661(C>T) in IGF-1 gene were found in Karan Fries cows, however rs29004509 (C>T) had positive correlation (p<0.05) with milk yield. The genetic variants observed in exon 3 region of *leptin* gene and their association with milk yield traits revealed the importance of CT genotype, which had been useful for genetic improvement of Karan Fries cow for milk production traits and can also be utilized as a potential genetic marker to select appropriate animals.

## 1. INTRODUCTION

In the current era of genomic informatics, attempts are increasingly underway to find certain meaningful associations in the genome that can be exploited to select animals for favorable production traits to attain rapid genetic gain in short time periods (Kiyici *et al*., 2019). Insulin-like growth factor-1 (IGF-1) gene is located on chromosome 5, consists of 4 exons spanning 66191602-66264083 bp length (Laron, 2001). IGF-1, a small peptide of 70 amino acids with a molecular weight of 7649 Dalton, is also considered to play an important physiological role in growth, development, metabolism and lactation in bovines (Mehmannavaz *et al*., 2010). IGF-1 is believed to be one of the main mediators of the effects of energy balance on the reproductive performance of the dairy cows after calving. Its concentrations are highly associated to postpartum energy balance (Mehmannavaz *et al*., 2010), follicular growth and resumption of ovarian cyclicity (Mullen *et al*., 2011; Lynch *et al*., 2010). SNPs, IGF1i3 A-G, IGF1i6 A-G, IGF1i7 A-G, rs29012855 A-G of IGF-1 have association (P < 0.05) with functional survival milk protein, fat yield, somatic cell score, carcass conformation and BCS in cows (Mullen *et al*., 2011). SNP/SnaBI in IGF-1 with milk production traits showed no significant effects within first 4 months of lactation but a significant association with interval to commencement of luteal activity in Holstein Friesian dairy cows have been reported (Nicolini *et al*., 2013). SNPs, 89C/T, 98G/T and 167T/C, of IGF-1 are associated with milk production and constituent traits in buffaloes (Fatima *et al*., 2009). The leptin, an adipokine also plays a significant role in several physiological processes including regulation of energy balance, fertility, milk production and was indicated as a candidate gene for marker assisted selection for high yielding cows (Liefers *et al.*, 2005, Vohra *et al.*, 2011, Kononoff *et al.*, 2017). Leptin gene is located on chromosome 7, consists of 3 exons, spanning 1282412782-128257629 bp length. Only exons 2 and 3 are translated, expressing 167 amino acids from which 21 amino acids as signal peptide are removed to leave a 16.7 kilo Dalton protein of 146 amino acids, arranged as four anti-parallel helices (A, B, C and D), which are typical class 1 cytokines (Zhang *et al.*, 1997). Previously an association between A1457G and milk yield in British cows have been found (Banos *et al.*, 2008; Clempson *et al.*, 2011). However, there are contradictory reports regarding association of leptin polymorphisms with composition and quality of milk (Ferreira *et al.*, 2019). Despite the studies on polymorphisms of leptin with respect to milk yield, there is no available literature on the relationship of these two genes on the same aspect. The present investigation was undertaken with a hypothesis that polymorphisms at the leptin and IGF-1 gene locus might play a role in milk production. The associations between leptin and IGF-1 gene polymorphisms and milk yield will provide insight into the underlying mechanisms of leptin and IGF-1 gene with milk production, and results may be used in future breeding programs. This study was undertaken to evaluate the association of single nucleotide polymorphism in bovine leptin and IGF-1 gene with the milk production performance in crossbred cows.

## 2. MATERIALS AND METHODS

### 2.1. Experimental animals and DNA extraction

The present experiment was carried out in indicine and taurine crossbred (Karan Fries) lactating cows of II and III parity at NDRI, Karnal, situated at an altitude of 250 meter above mean sea level, latitude and longitude position being 29° 42” N and 79° 54” E respectively. Experiment was approved by the Institutional Animal Ethics Committee (IAEC) constituted as per the article no.13 of the CPCSEA rules, laid down by Govt. of India. Crossbred Karan Fries cows (n=160) were selected from the institute livestock research center of ICAR-NDRI, Karnal during the month of June. The cows were maintained in loose housing system and were fed as per the ICAR (2013) feeding standard, received green fodder and concentrate mixture as per the requirement in the ratio of 60:40. The cows were machine milked three times a day in the morning, noon and evening and the milk yields were recorded. Ten ml of blood was collected in EDTA coated vaccutainer tubes in morning and was stored at −20°C until DNA isolation. Genomic DNA was extracted from the blood samples using phenol-chloroform extraction method with minor modifications (Sambrook and Russell, 2001). The quality of DNA was checked by 1.5% agarose gel electrophoresis. Quality and quantity of DNA was also estimated by Biospec-nano spectrophotometer (Shimadzu co-operation, Japan). The ratio between OD_260_ and OD_280_ was observed for each sample. DNA sample with a ratio of 1.8 was further diluted to a final concentration of 30 ng/μl and was stored at −20°C for further analysis.

### 2.2. Polymerase chain reaction (PCR) primers and amplifications

The primer was designed based on the bovine leptin and IGF-1 gene sequence (NCBI GenBank AC_000162.1 and AC_000161.1) using Primer3 software (Table 1). The PCR reactions were carried with 25 μl total volume containing template DNA of 3 μl (30 ng/μl), 1.0 μl of forward and reverse primer, PCR Master Mix (2x) (Fermentas) of 12.5 μl, and 8.5 μl of water. Amplification was performed in a Thermal cycler (MJ research and BioRad, T100). The thermal cycling conditions involved an initial denaturation at 95°C for 3 min, followed by 35 cycles with initial denaturation at 95°C for 30s, annealing temperature of 57.9°C and 57.3°C for 30 sec respectively, extension at 72°C for 1 minute. PCR program for both the primers was similar and the PCR products were detected by electrophoresis on 2% agarose gel stained with ethidium bromide.

**Table 1.**
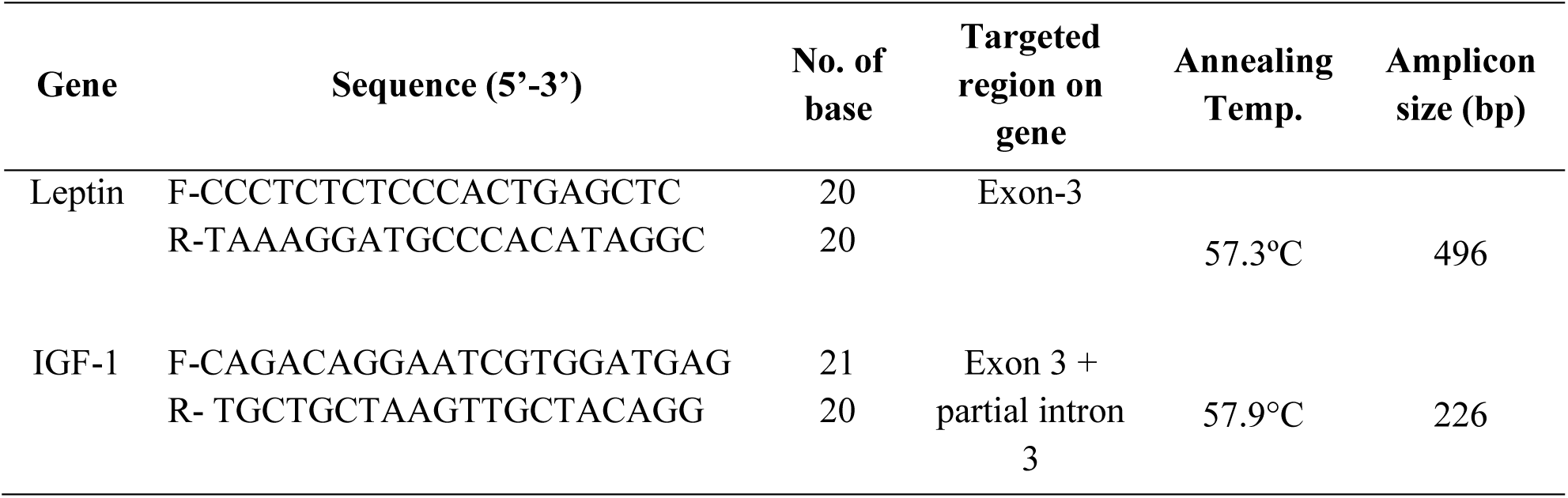
Primer sequences, targeted region, and amplicon sizes of bovine leptin and IGF-1 gene

The gene and genotypic frequencies of different genotypes were estimated by standard procedure POPGENE version 1.32 (University of Alberta, Canada) (Yeh *et al.*, 1999). The association of SNP genotypes with milk yield was analyzed using General Linear model (SAS Version 9.2) as follows: 

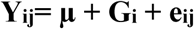

where,

Y_ij_ = Adjusted value of milk yield of j^th^ animal of i^th^ genotype

µ = Overall mean

G_i_ = Effect of i^th^ Genotypes

e_ij_ = Residual error NID (0, σ^2^_e_)

## 3. RESULTS

### 3.1. Analysis of sequence data association of milk yield traits with SNPs

The PCR product with the amplicon size of 496 and 226 bp was successfully amplified, covering exon 3 region of *leptin* gene and exon 3 + partial intron 3 of *IGF-1* gene in Karan Fries cows, respectively (Figure 1, 2). The final sequence of the contig for Karan Fries cows were deduced from the raw sequences by using BioEdit software. Clustal Omega software, with a reference sequence of *Bos taurus* (NCBI GenBank AC_000162.1 and AC_000161.1) was used for determining the polymorphism in exon 3 of Karan Fries cow. The comparison of nucleotide sequences of exon 3 of leptin and exon 3 + partial intron 3 of IGF-1 gene with that of reference sequence of *Bos taurus* by Clustal Omega multiple alignments revealed 6 mutations including 5 transitions and 1 transversion. Four SNPs were found at positions rs29004508 (C>T), rs29004509 (C>T), rs29004510 (T>C), rs29004511 (T>C) in Leptin gene and two SNPs at positions rs133251968 (C>A), rs137289661(C>T) in IGF-1 gene as compared to *Bos taurus* (Ref. Seq. AC_000161.1 and AC_000162.1; Table 2 and Figure 3). The association of leptin and IGF-1 gene with milk yield elucidated SNP rs29004509 amplified by the primer 1 of leptin gene correspond to two genotypes CT and CC (Table 3). However, CT genotype of SNP rs29004509 had positive effect (P<0.05) on milk production with mean of 3827.06 ± 145.394 (Table 4). Rest of the identified SNPs like rs29004508 (C>T), rs29004510 (T>C), rs29004511 (T>C), rs133251968 (C>A), rs137289661(C>T) were not correlated with milk yield. The genotype and allelic frequencies of leptin and IGF-1 gene are represented in table 3, respectively.

**Table 2.** Nucleotide changes in leptin and IGF-1 gene in Karan Fries cattle as compared to *Bos taurus* (NCBI Gene ID AC_000161.1 & AC_000162.1, respectively)

| Gene | SNP | Location | Nucleotide Change | Type of Variation | Amino Acid Substitution |
| --- | --- | --- | --- | --- | --- |
| Leptin | rs29004508 | Exon 3 | (C>T) | Transition | p.80A>V |
|  | rs29004509 |  | (C>T) | Transition | - |
|  | rs29004510 |  | (T>C) | Transition | - |
|  | rs29004511 |  | (T>C) | Transition | - |
| IGF-I | rs133251968 | Intron 3 | (C>A) | Transversion | - |
|  | rs137289661 |  | (C>T) | Transition | - |

**Table 3.**
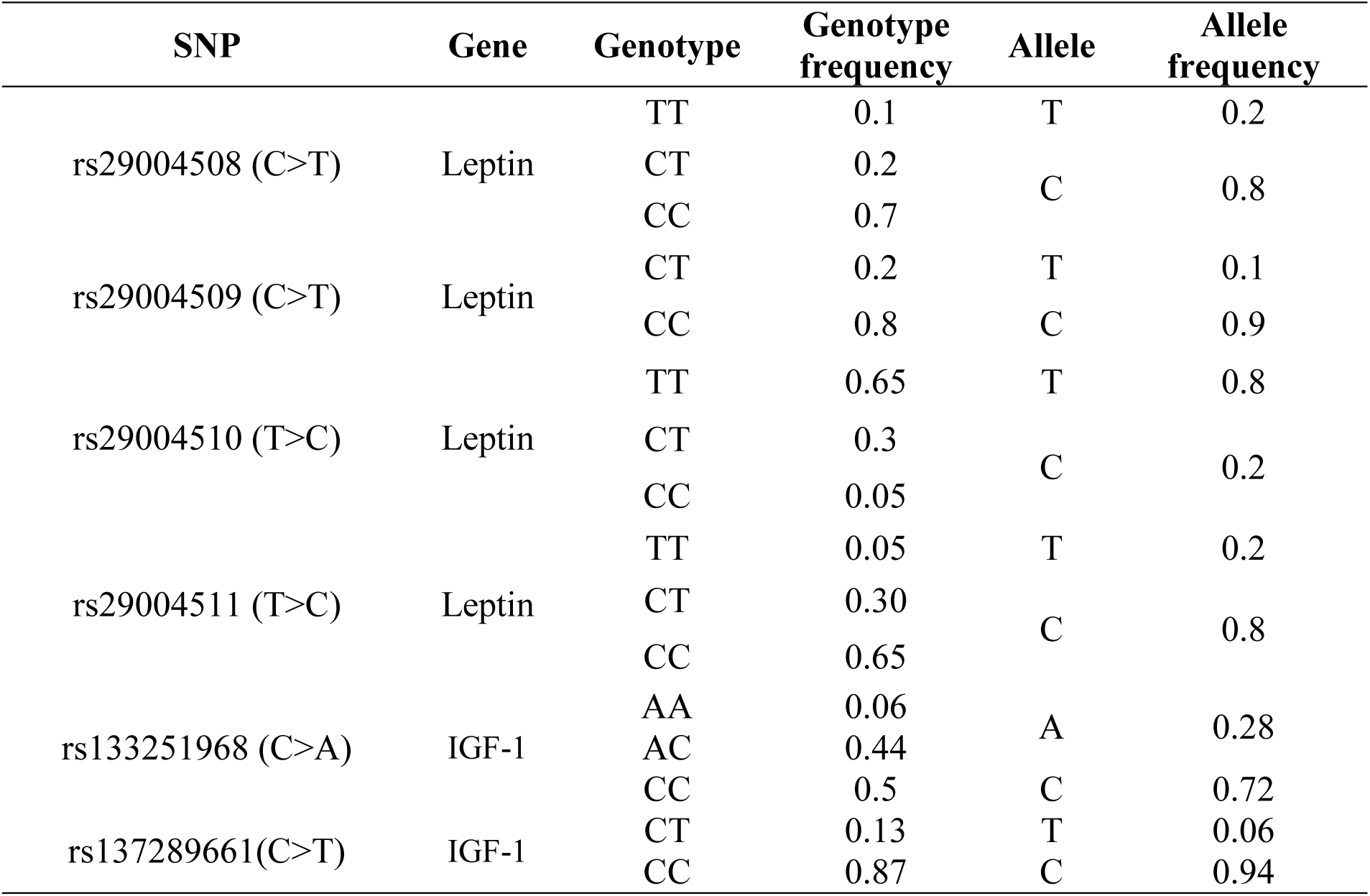
Genotypic and allelic frequencies of leptin and IGF-1 gene using sequencing in Karan Fries cattle

**Table 4.** Effect of polymorphism of leptin and IGF-1 gene on average milk yield in different genotypes

| SNP | Genotype | Average milk yield (Kg) | p-value |
| --- | --- | --- | --- |
| rs29004508 (C>T) | TT | 3587.19 ± 153.04 | 0.899 |
|  | CT | 3882.20 ± 290.08 |  |
|  | CC | 3746.27 ± 289.59 |  |
| rs29004509 (C>T) | CT | 3827.06<sup>a</sup> ± 145.39 | 0.021 |
|  | CC | 3280.22<sup>b</sup> ± 242.99 |  |
| rs29004510 (T>C) | TT | 3642.67 ± 528.71 | 0.162 |
|  | CT | 3599.12 ± 193.18 |  |
|  | CC | 3753.37 ± 169.62 |  |
| rs29004511 (T>C) | TT | 3753.33 ± 169.62 | 0.162 |
|  | CT | 3599.12 ± 193.18 |  |
|  | CC | 3642.67 ± 528.71 |  |
| rs133251968 (C>A) | AA | 3836.63 ± 201.80 | 0.521 |
|  | AC | 3211.43 ± 277.11 |  |
|  | CC | 3073.50 ± 465.40 |  |
| rs137289661(C>T) | CT | 3551.92 ± 180.98 | 0.453 |
|  | CC | 3539.57 ± 296.18 |  |

Values with different superscripts <sup>a, b</sup> differ between genotypes along with their average milk (P<0.05)

**Figure 1:**
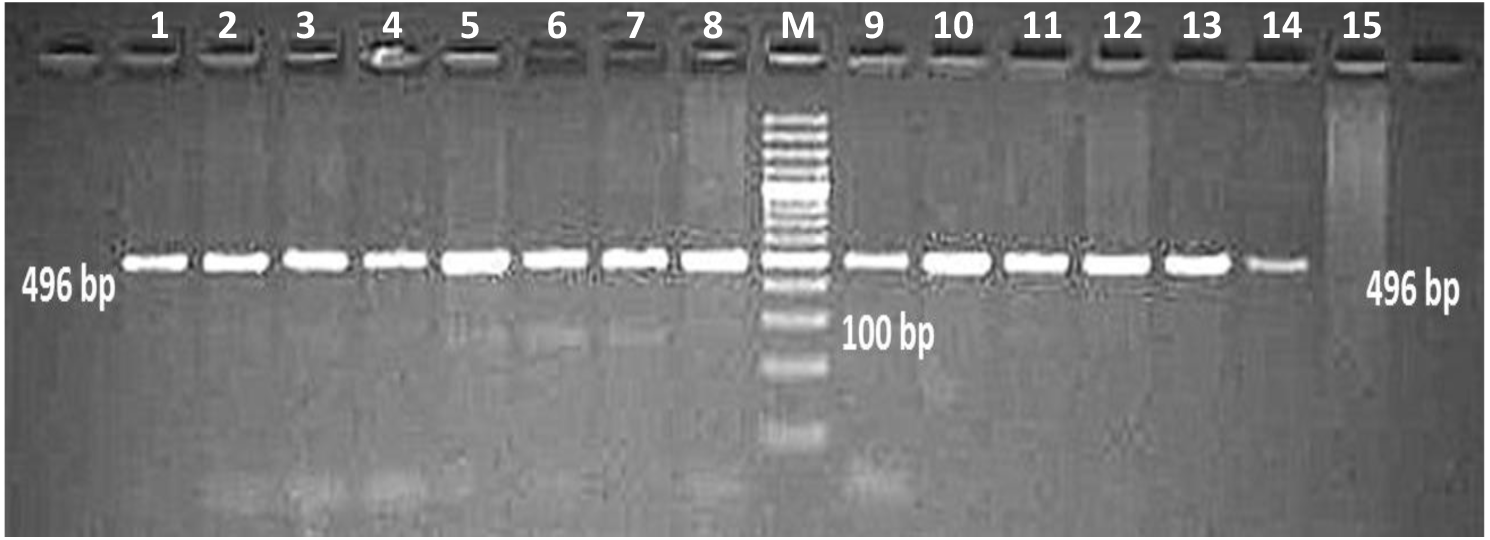
PCR amplification of Exon 3 of leptin gene in Karan Fries cow (M=100bp) Lanes 1-15 = 496 bp, Lane M = 100 bp

**Figure 2:**
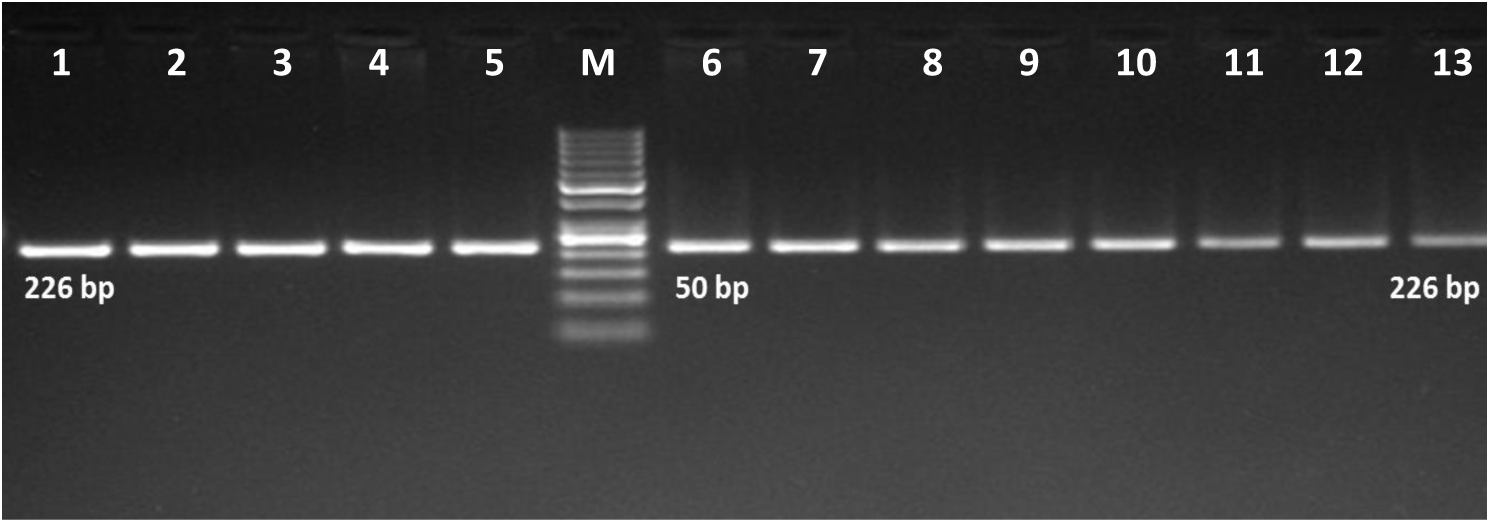
PCR amplification of Exon 3 + some part of intron 3 of IGF-1 gene in Karan Fries cow (M=50bp) Lanes 1-15 = 226 bp Lane M = 50 bp

**Figure 3.**
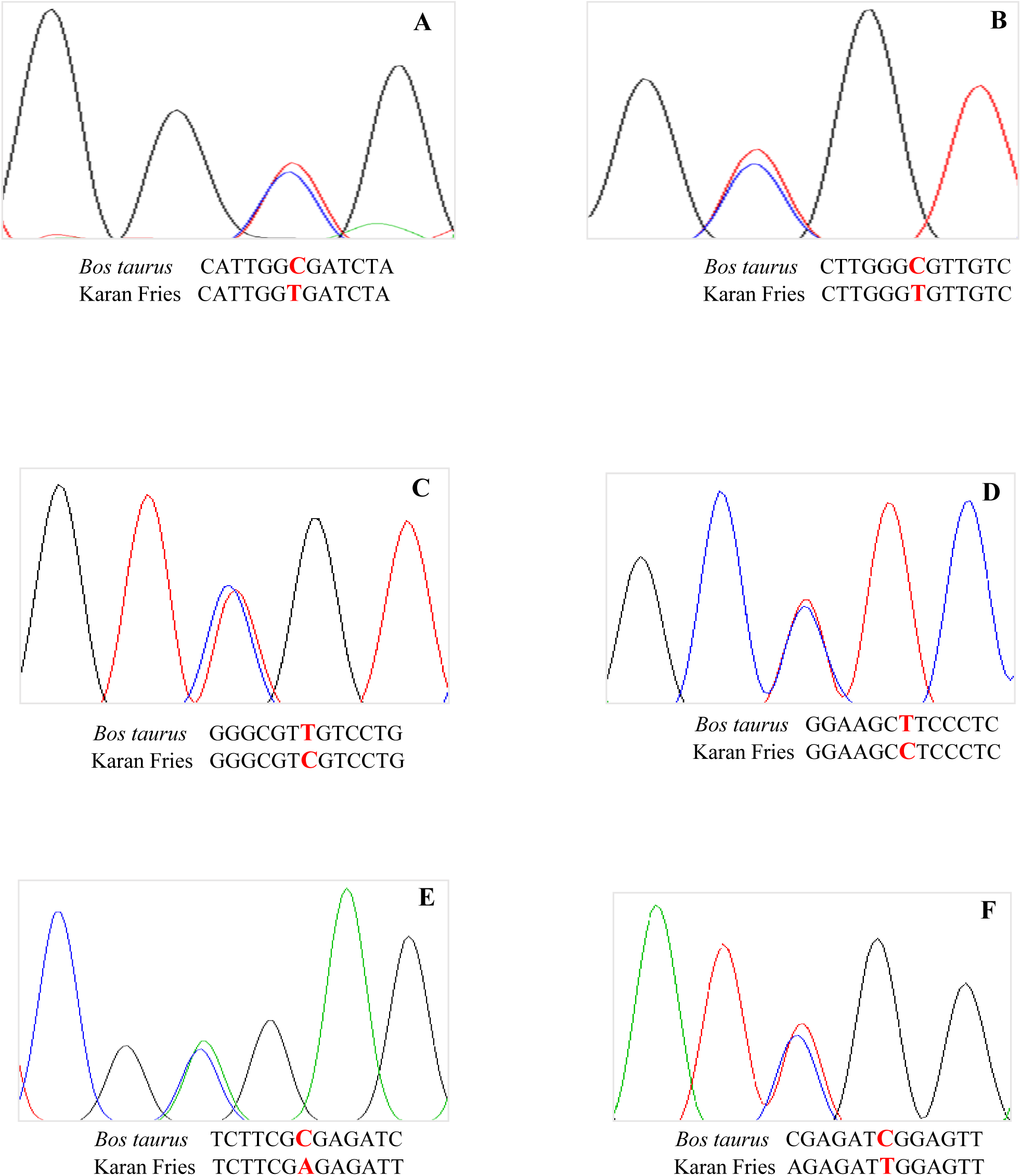
Chromatogram and Clustal Omega alignment showing variation at position (A) rs29004508 (C>T), (B) rs29004509 (C>T), (C) rs29004510 (T>C), (D) rs29004511 (T>C) for Leptin gene and (E) rs133251968 (C>A), (F) rs137289661(C>T) for IGF-1 gene in Karan Fries cattle.

## 4. DISCUSSION

The present study revealed presence of four SNPs at positions rs29004508 (C>T), rs29004509 (C>T), rs29004510 (T>C), rs29004511 (T>C) in Leptin gene and two SNPs at positions rs133251968 (C>A), rs137289661(C>T) in IGF-1 gene in crossbred cows in comparison to *Bos taurus*. Only rs29004509 (C>T) showed positive correlation (p<0.05) with the milk production in Karan Fries cows. Similar to our results, Nobari *et al.* (2011) reported that AB genotype had a significant effect on milk production, days open and milking days compared to other genotypes in Brown Swiss cows and Liefers *et al.* (2002) reported that AB genotype had higher milk production in Holstein heifers. Association between polymorphism of leptin gene and milk production have been reported in several studies, leptin SNP A1457G association with milk production was found significant in cattle (Banos *et al.*, 2008) and high producing cattle (Clempson *et al.*, 2011). The HphI-RFLP locus had significant effect on average daily milk yield, while, Kpn2I-RFLP had significant effect on first lactation milk yield and average daily milk yield (Choudhary *et al.*, 2019). An association of single nucleotide polymorphisms (SNP) in the leptin gene with production efficiency as well as milk protein and milk yield has been reported (Barendse *et al.*, 2004). The association of leptin gene polymorphism with growth traits and reproduction traits has been reported in crossbred and indigenous cattle (Choudhary, 2004). GH-TaqI, LEP-Sau3AI and MYF5-TaqI polymorphisms of the genes investigated had significant effects, especially on the 305-day milk yield of Holstein cows (Sahin and Akyuz, 2017, Kiyici *et al.*, 2019). Trakovicka, (2013) reported that the SNP LEP/Sau3AI significantly influence milk, protein and fat yield (P<0.05) in cows. Similarly polymorphism with respect to leptin gene was highly significant with milk yield and milk composition traits in Polish Black and White cows (Polish Friesian) (Flisikowsk *et al.*, 2004). Lende *et al.*, (2005) assessed that the Leptin SNP have association with milk yield, milk composition and DMI in dairy cows. Many SNPs were detected on the promoter region as well as exon regions of the leptin gene, and were found to be highly associated with different milk traits. Karan Fries cattle with TT genotype showed significantly higher 305 days milk yield as compared to cattle with CC genotype (Vohra *et al.*, 2011). Other reports showed that C/BspEI/T and C/HphI/T poly-morphisms of leptin gene is associated with milk protein percentage, whereas the C/HphI/T locus of leptin is significantly associated with Solid non fat (SNF) percentage (Singh *et al.*, 2014). Contrary to our findings, Ferreira *et al.* (2019) reported that polymorphisms leptin do not influence the composition and quality of milk from 1/2, 3/4 and 7/8 Holstein x Guzera cows kept in a hot climate.

Kulig and Kmiec (2009) evaluated that the selection for the A59V CC and CT animals might contribute to enhance in milk yield as well as fat and protein yields in Jersey cattle. Contrary to our findings, Mullen *et al.* (2011) reported that SNPs, IGF1i3 A-G, IGF1i6 A-G, IGF1i7 A-G, rs29012855 A-G, of IGF-1 are associated (P < 0.05) with functional survival and chest width, milk protein and fat yield, milk fat concentration, somatic cell score, carcass conformation and fat. In another study, (Lynch *et al.*, 2010) SNPs, IGFi1A-T, IGFi2C-T, IGFi3G-A, in IGF-1 were associated with milk production traits and BCS; and are in the agreement with our findings. Other studies reported that the polymorphism of bovine IGF-1 gene in exon 4 were associated with production traits in Bali cattle (Maskur *et al*., 2012) but polymorphism in the intron 4 of IGF-1 gene was not associated with production traits in mixed population of Charolais and Beef master cattle (Reyna *et al*., 2010).

It has been found that the SNP IGF-1/SnaBI with milk production traits have no significant effects on either milk yield, fat corrected milk or Total solids yield within first 4 months of lactation (Nicolini *et al.*, 2013). In dairy cattle, some studies have analyzed the association of this SNP (IGF-1/SnaBI) with milk production traits (Hines *et al.*, 1998; Siadkowska *et al.*, 2006; Bonakdar *et al.*, 2010; Mehmannavaz *et al.*, 2010; Mullen *et al.*, 2011; Ruprechter *et al.*, 2011). Mehmannavaz *et al.* (2010) proved a significant effect of the SNP *IGF-1*/*Sna*BI on estimated breeding values (EBV) for milk production traits in Iranian Holstein bulls, as animals with AB genotype had higher EBV for milk and fat yields than homozygous genotypes. The interval from calving to commencement of luteal activity postpartum (CLA) has been suggested as a suitable selection criteria for fertility, as early CLA is an important factor for a new pregnancy after calving, it presents higher heritability values (16 to 25%) than traditional fertility traits, and it is genetically favorably correlated with traditional fertility traits (Darwash *et al.*, 1997; Petersson *et al.*, 2007; Nicolini *et al.*, 2013). SNPs, 89C/T, 98G/T and 167T/C, of IGF-1 are reported to be significantly associated with milk production and constituent traits in buffaloes (Fatima *et al.*, 2009).

## CONCLUSIONS

In this study novel SNPs were detected in indicine and taurine crossbred cows. SNP at position rs29004509 (C>T) in Leptin gene had positive correlation (p<0.05) with milk yield. The genetic variants observed in exon 3 region of *leptin* gene and their association with milk yield traits revealed the importance of CT genotype, which can be utilized as a potential genetic marker to select elite cows for genetic improvement in future.

## ACKNOWLEDGMENTS

The authors are thankful to the Director, National Dairy Research Institute, Karnal Haryana for providing the necessary facilities to carry out this experiment. This work was supported by Board of Research in Nuclear Science (BRNS) Mumbai, India under project no. 2013/35/48/2013-BRNS/RTAC.

## AUTHOR’S CONTRIBUTIONS

MR, MS and AV designed the study. MR and ST provided the data and performed the analysis. MR, MS and AV drafted the manuscript. MR, MS, ST and AV contributed to the interpretation of results, the discussion and commented on the manuscript. All authors read and approved the manuscript.

## ORCID

http://orcid.org/0000-0003-3541-7239

